# Shifts in honeybee worker metabolism immediately post-eclosion

**DOI:** 10.1101/2024.12.01.622772

**Authors:** Gilles Verbinnen, Mikkel Roald-Arbøl, Jeremy Edward Niven, Elizabeth Nicholls

## Abstract

- The metabolic rate of an organism is intrinsically linked to key traits such as reproductive output and lifespan. While the drivers of individual differences in metabolic rate are poorly understood, previous research in insects has shown that metabolic rate can change substantially in the initial hours and days post-eclosion as adults.
- Here we repeatedly measured the resting and active metabolic rate of individual adult honeybees (*Apis mellifera*) for up to 48 hours from the time of eclosion. We combined flow-through respirometry with automated behaviour tracking, permitting us to obtain ‘active’ (AMR) and true ‘resting’ metabolic rate (RMR) from freely moving animals. We compared these recordings to the more conventional approach of obtaining resting metabolic rate by restraining animals.
- Both active and resting metabolic rates and mass-specific metabolic rates increased significantly in the first 48-hours post-eclosion, whereas metabolic scope did not change. Mass-specific water loss was highest in active bees and changed non-linearly with time post-eclosion, increasing in the first 24 hours before decreasing again. A similar quadratic relationship with time was also observed for bees’ movement speed. Speed- and mass-specific metabolic rate and scope increased with time post-emergence, whereas speed- and mass-specific water loss did not.
- The metabolic rate of restrained bees was consistently significantly higher than the true RMR at all time points, likely due to the stress associated with being restrained. Therefore, we recommend future studies of insect resting metabolic rates avoid restraining organisms to restrict movement and consider employing behaviour tracking as a means to extract metabolic rate data from periods of true rest.
- This study provides important insights into the previously overlooked changes in metabolism exhibited by newly emerged honeybee workers. The high mortality rate beyond 48 hours, coupled with significant changes in metabolic rates, body mass, and water loss, underscores the importance of this early post-eclosion period for survival and metabolic stabilization.

## Introduction

An organism’s resting metabolic rate (RMR) is a measure of the energy consumed in maintenance, though in many animals, including insects, it is measured when the animal is stationary and so includes muscular activity needed to maintain posture. In contrast, active metabolic rate (AMR) includes the energy consumed through movement and is dominated by skeletal muscle (Weibel, 2002; Weibel *et al*., 2004) and is consequently, dependent on the activity undertaken. The ratio of the AMR to RMR (AMR/RMR) is the metabolic scope (Peterson *et al*., 1990). Metabolic rates are dynamic and vary according to internal factors such as developmental stage as well as environmental conditions such as temperature, nutrition and/or food availability (Cortés et al., 2015; Harrison et al., 2012; Norin & Malte, 2011; White et al., 2013). Both RMR and AMR show considerable intra-specific variation and have been linked to life history traits such as lifespan and reproduction (Burton et al., 2011; Glazier, 2005; Kasumovic & Seebacher, 2013; Niitepõld & Hanski, 2013; Pettersen et al., 2018; Speakman, 2005). While the drivers of intra-specific variation in metabolic rate are still quite poorly understood (Konarzewski & Ksiazek, 2013), the optimal metabolic rate for an organism is likely context-dependent (Burton et al., 2011; Norin & Metcalfe, 2019; Zeng et al., 2017).

Holometabolous insects undergo substantial changes in external and internal morphology and physiology throughout their development from larva to pupa to adult (reviewed in Chapman 2013). However, even in the immediate hours and days post-eclosion adult holometabolous insects continue to undergo morphological, physiological and behavioural maturation processes that might be expected to influence both RMR and AMR. Many such processes, including cuticular hardening and flight muscle development are energetically demanding, therefore it is unsurprising that in various insects metabolic rates are highest during the initial hours post-eclosion and gradually stabilize as the insects complete their maturation (Harrison & Roberts, 2000; Piiroinen et al., 2010; Terblanche et al., 2004). In the Colorado potato beetle (*Leptinotarsa decemlineata)*, for example, RMR increases significantly post-eclosion, peaking at 48 hours before declining again (Piiroinen et al., 2010). Increases in RMR between 24-196 hours post-eclosion have also been shown in tsetse flies (*Glossina pallidipes*) (Terblanche et al., 2004). Water regulation during this period is critical because newly emerged insects often face challenges with water loss due to incomplete maturation of the cuticular exoskeleton (Gibbs et al., 2003; Gibbs & Markow, 2001). Water loss can have a direct impact on metabolic processes because dehydration can limit an insect’s ability to maintain necessary physiological functions, further influencing their metabolic rate (Hadley, 1994).

Honeybee (*Apis mellifera* L.) workers offer an interesting system for exploring the drivers of intra-specific variation in metabolic rates, given it is possible to examine the impact of variation in the genetic background by studying genetically identical workers from the same colony (Harrison & Fewell, 2002). In the first few days post-eclosion, adult honeybee workers undergo significant physiological and behavioural changes crucial for their development and future roles in the hive, that might be expected to drive rapid changes in metabolism. One of the most important is the growth of their hypopharyngeal glands, that produce royal jelly enabling workers to start feeding larvae and the queen (Crailsheim & Stolberg, 1989; Huang & Robinson, 1996). At the same time, juvenile hormone (JH) levels begin to increase, influencing the maturation of various organs and the bees’ eventual shift from in-hive tasks, such as nursing, to outside roles like foraging (Jassim et al., 2000; Sullivan et al., 2003; Wegener et al., 2009). Flight muscle development also begins over this period. Stabentheiner et al. (2010) found that newly emerged bees (0–2 days old) have limited thermoregulatory capacity, remaining ectothermic until their flight muscles develop at around 2 days old, at which point they can thermoregulate. The switch from earliest in-hive task of cleaning brood cells ready for queen to lay eggs, which involves walking across the comb (Ribbands, 1952; Robinson, Page & Huang 1994), to the energetically demanding tasks of flying and foraging is thought to be driven, at least in part, by developmental shifts in metabolism (Harrison & Roberts, 2000; Schippers et al., 2006; Scofield & Amdam, 2024; Toth & Robinson, 2005). Yet, studies of metabolic rate in honeybees at different life stages have found no significant differences between the RMR of younger in-hive bees (7-10 days old) and older foragers (Kovac et al., 2007; Stabentheiner et al., 2003). Whilst Schippers et al. (2010) observed an initial increase in flight metabolic rate (AMR) of 68% between one day old and 3-4 day old in-hive bees that plateaus and doesn’t increase again until the onset of foraging.

Few studies have explored the changes that may occur in the hours and days immediately post-eclosion, when honeybees’ in-hive behaviour is dominated by walking and local limb movements, meaning that rapid and/or transient increases in RMR described in other insects may have been missed. Allen (1959) did record a small number of restrained (caged) honeybee workers immediately post-eclosion but did not measure the metabolic rates of the same individuals at different time points. Indeed, honeybee workers are often restrained in studies of metabolic rates (Karise et al., 2016; Nicholls et al., 2021) because of their high activity levels (Kovac et al., 2007) that make it challenging to measure RMR. Yet, the metabolic rate of restrained insects does not necessarily provide an accurate estimate of RMR (Perl & Niven, 2018). One means of avoiding restraint is to record trials with video, tracking individuals to identify periods of activity and spontaneous rest. Accurately tracking individuals manually can be time consuming, particularly for long recordings. New developments in the field of automated behaviour tracking and the accessibility of open-source software such as ‘TRex’ (Walter & Couzin 2021) and ‘Animal TA’ (Chiara et al., 2023), mean that measuring the metabolic rates of unrestrained animals and using tracking to partition behaviour into periods of rest and activity is feasible.

To our knowledge, studies have not measured RMR and AMR in the *same* individual honeybee workers within the first few days post-eclosion, permitting examination of the maturation of morphology, physiological and behavioural changes. Here we repeatedly measured body mass, RMR, AMR, metabolic scope, water loss and movement speed from individual unrestrained adult honeybee workers’ for up to 96 hours post-eclosion. RMR was measured during periods in which they were stationary and AMR from periods in which they were walking, corresponding to their in-hive behaviour immediately post-eclosion. To achieve this, we combined flow-through respirometry with automated behaviour tracking. We also compared these recordings to the metabolic rate in restrained animals, to test whether the same individuals differed between methods.

## Methods

### Bee rearing

Capped frames of honeybee (*Apis mellifera* L.) brood were obtained from four hives kept on campus grounds at the University of Sussex, UK. Frames were placed inside a frame holder and maintained in an incubator held at 35°C and 60% relative humidity (RH). Frames were checked a minimum of four times per day for the eclosion of new workers, and their metabolic rate (see below) was individually measured on the day of eclosion (0-4 hours old, referred to as 0h hereafter) and every 24 hours for the next 5 days (24, 48, 72 and 96 hours post-eclosion).

After the first recording, bees were kept in individual queen roller cages (Thorne, UK) in an incubator (35°C, 60% RH), with *ad libitum* access to deionised water, delivered via damp paper towels placed under the roller cages. Each day, immediately after a recording, bees were individually fed via a syringe with 50 mg of 30% sucrose solution (w/w), that was supplemented with a protein and vitamin-rich pollen supplement Vitafeed nutri (Vita Beehealth, UK) at a rate of 5g/kg of sucrose solution (14% crude protein). Survival was monitored daily, and any deceased bees were frozen at -20°C.

### Measurement of metabolic rate and water loss

To test whether adult honeybee workers’ RMR, AMR and water loss changes in the days post-eclosion, we used flow-through respirometry, with CO_2_ production over time as a measure of metabolic rate. We also measured H_2_O production over the same period. Each bee was tested at a similar time each day to reduce the potential influence of circadian rhythms. Recordings of individual bees were made under two consecutive conditions within the same daily trial. For the first 15 minutes of a recording, bees were restrained using a small cylinder of metal mesh to permit gas exchange but prevent bee movement. In the second half of the recording (15 minutes) the metal mesh was removed, and bees were unrestrained and able to move freely in the test chamber. Recordings were always performed in this order and each recording lasted 30 minutes in total. The room temperature was maintained at 20.5°C (±2.5°C) and all tests were performed in the dark to resemble conditions within the hive. Bees were weighed before and after each recording. After weighing, an individual bee was placed inside a chamber, adapted from a 15 mL centrifuge tube (Corning Falcon, USA), with custom-designed 3D-printed closures at each end. Air scrubbed of CO_2_ and H_2_O was pumped through the chamber at a consistent rate of 200 mL min^-1^ regulated by mass flow controller (GFC17; Aalborg, NY, USA) positioned immediately prior to the test and identical, empty reference chamber. Air from the chambers then passed through an infrared CO_2_-H_2_O analyser (Li7000, Li-Cor Biosciences, Lincoln, Nebraska, USA). During tests with restrained bees, an identical piece of metal mesh was placed in the reference chamber. CO_2_ and H_2_O production were measured in differential mode (Nicholls et al., 2017; Perl & Niven, 2018) (Fig. 1).

**Figure 1.**
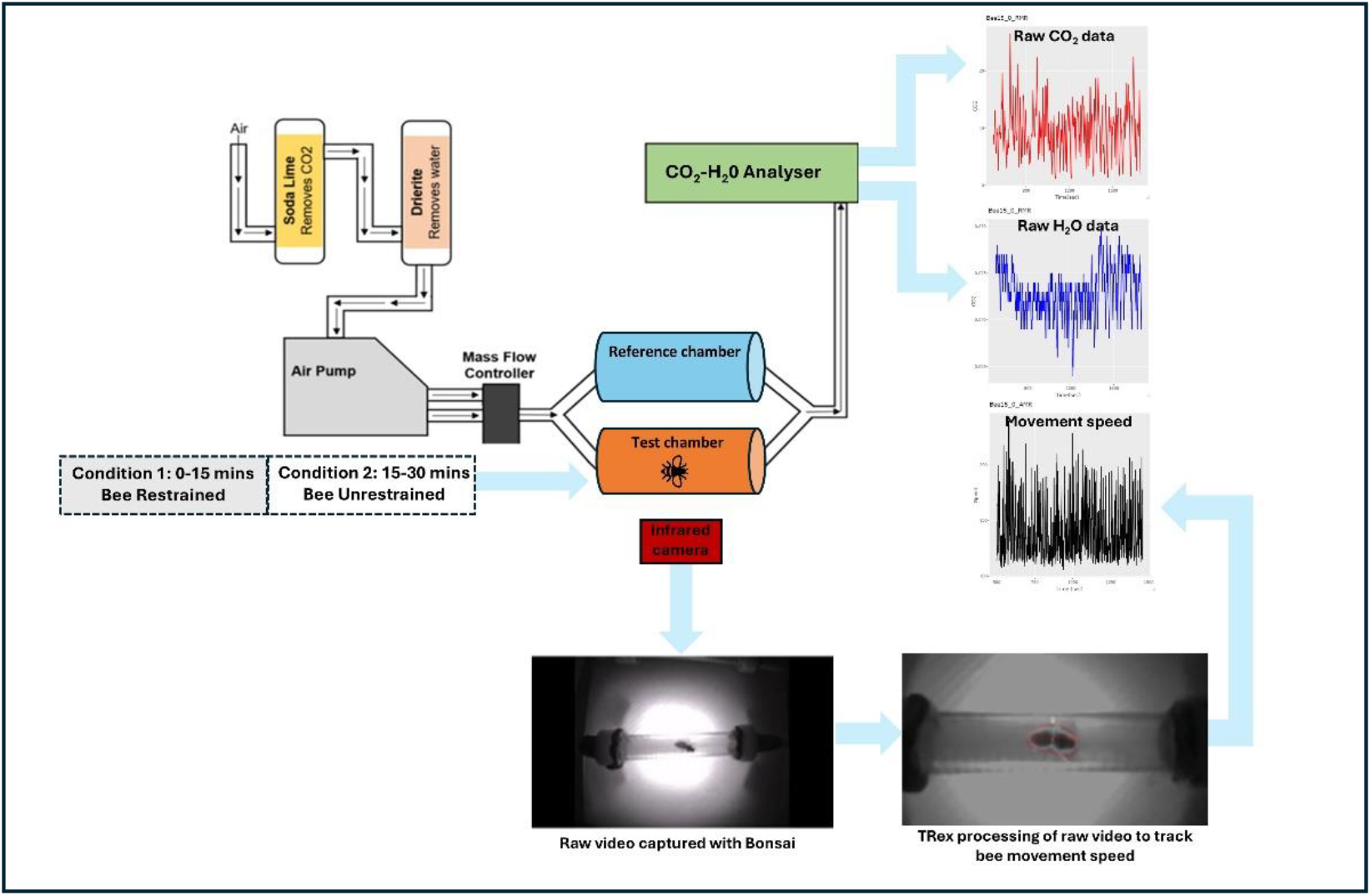
Schematic overview of the combined respirometry and automated-behaviour tracking set up used to extract data on bees’ metabolic rates, rates of water loss and movement speed, which was subsequently used to partition the behaviour of unrestrained bees into periods of activity and rest (defined as a period of at least three minutes where the speed was below 0.03 cm/s).

### Tracking bee behaviour

To identify periods of activity and rest during the unrestrained metabolic rate recording, the behaviour of unrestrained bees was filmed from above at a rate of 30 frames per second (fps) using an infrared camera suspended from a scaffold (MakerBeam B.V., Utrecht, Netherlands). The respirometry chamber was positioned on top of a 3D-printed platform and illuminated from below using infrared LEDs; light was diffused using photographic fabric. Raw video footage was captured with Bonsai (Lopes et al., 2015) and then processed with the automatic tracking software TRex (v1.1.8_2) (Walter & Couzin 2021) to extract the position of the bees (see Supplementary Materials).

### Exclusion of bees beyond 48 hours post-eclosion

Despite feeding bees and keeping them in a temperature and humidity-controlled incubator, mortality beyond 48 hours post-eclosion was over 70% (Fig. S1), leaving just 11 bees to test at 72 hours and 10 bees at 96 hours. Given these small sample sizes, we excluded metabolic and behavioural data from the 72- and 96-hour recordings from all analyses. Thereby minimising the risk of drawing conclusions from the behaviour and physiology of a small cohort of bees unlikely to be representative of newly emerged workers.

### Data analysis

All data analysis was performed in R (v. 4.4, R Core Team, 2022. https://www.R-project.org) within RStudio (v. 4.2.1) (see Supplementary Materials). Bees’ movement speed was smoothed with a rolling median filter over 30 frames. The recordings of unrestrained bees were then divided into active and resting periods based on speed. Resting was defined as a period of at least three minutes where the speed was below 0.03 cm/s. This resulted in 85 restrained recordings (0h= 37; 24h = 31; 48h =17), 35 unrestrained resting recordings (0h= 18; 24h = 10; 48h =7) and 82 active recordings (0h= 37; 24h = 29; 48h =16). For resting recordings, we removed the first and last 10 seconds of the recording to ensure transitions between activity and rest were not included.

Volumes of CO_2_ and H_2_O were temperature normalised to 20.5°C using the Q_10_ correction. To calculate the rate of CO_2_ production per bee (μl/hour), the volume of CO_2_ (ppm) was converted to CO_2_ fraction and multiplied by the flow rate (200 mL min^-1^). H_2_O production was converted to a rate of ng/hour. The CO_2_, H_2_O and speed recordings were then integrated using the ‘*trapz*’ function from the “*pracma*” package (Borchers 2023).

To visualise how bees’ survival changed with time post-eclosion we plotted a Kaplan-Meier survival curve using the ‘*survfit*’ function from the “*survival*” package (Therneau et al. 2024). To investigate the relationship between time and bee activity, we employed a generalized linear mixed model (GLMM) with a beta distribution using the “glmmTMB” package (Brooks et al. 2024). The beta distribution was selected as the response variable, activity, represents a proportion (time spent active), bounded between 0 and 1. However, since beta regression requires strictly positive values between 0 and 1, we adjusted any activity values of exactly 0 or 1 to 0.0001 and 0.9999, respectively, to ensure compatibility with the model. The model included time as a fixed effect to account for the influence of time on activity levels. In this and all other models, bee identity (bee_id) was included as a random effect to account for repeated recordings from the same individuals over time (0, 24 and 48 hours) and between different recording conditions (restrained *vs*. unrestrained).

Mixed effect models were used to compare the metabolic rate between unrestrained and restrained bees, and active and resting bees over time post-eclosion, and the fixed effects of time post-eclosion and activity level (active vs. resting) on water loss, mass-specific metabolic rate and mass-specific water loss. Mixed effect models were also used to test for changes in body mass, movement speed and mass and speed-specific metabolic rate, water loss and scope of active bees over time. Where relationships between the dependent variable and time appeared quadratic, we included the term (time^2^). Adequacy of the model fits was assessed from diagnostic plots of the standardised residuals (quantile–quantile and residuals over fitted). Non-significant interactions were dropped from the model. Estimated marginal means (emm) and pairwise comparisons were obtained using the “*lsmeans*” package (Length & Lenth 2018) and the *p*-value was adjusted for multiple comparisons with the Tukey method. All plots were made using the “ggplot2” package (Wickham 2016).

## Results

### Unrestrained bees show consistently high levels of activity over time

Bees typically spent more than three-quarters of their time within the chamber active (Fig. 2a; Table S1, Median + inter-quartile range (IQR) 0h= 1 (1-0.68) n= 37; 24h = 1 (1-0.73) n= 29; 48h = 1 (1-0.54) n=16) and this level of activity was consistent over time (GLMM with beta-distribution χ^2^= 0.800, df= 2, p=0.671).

**Figure 2.**
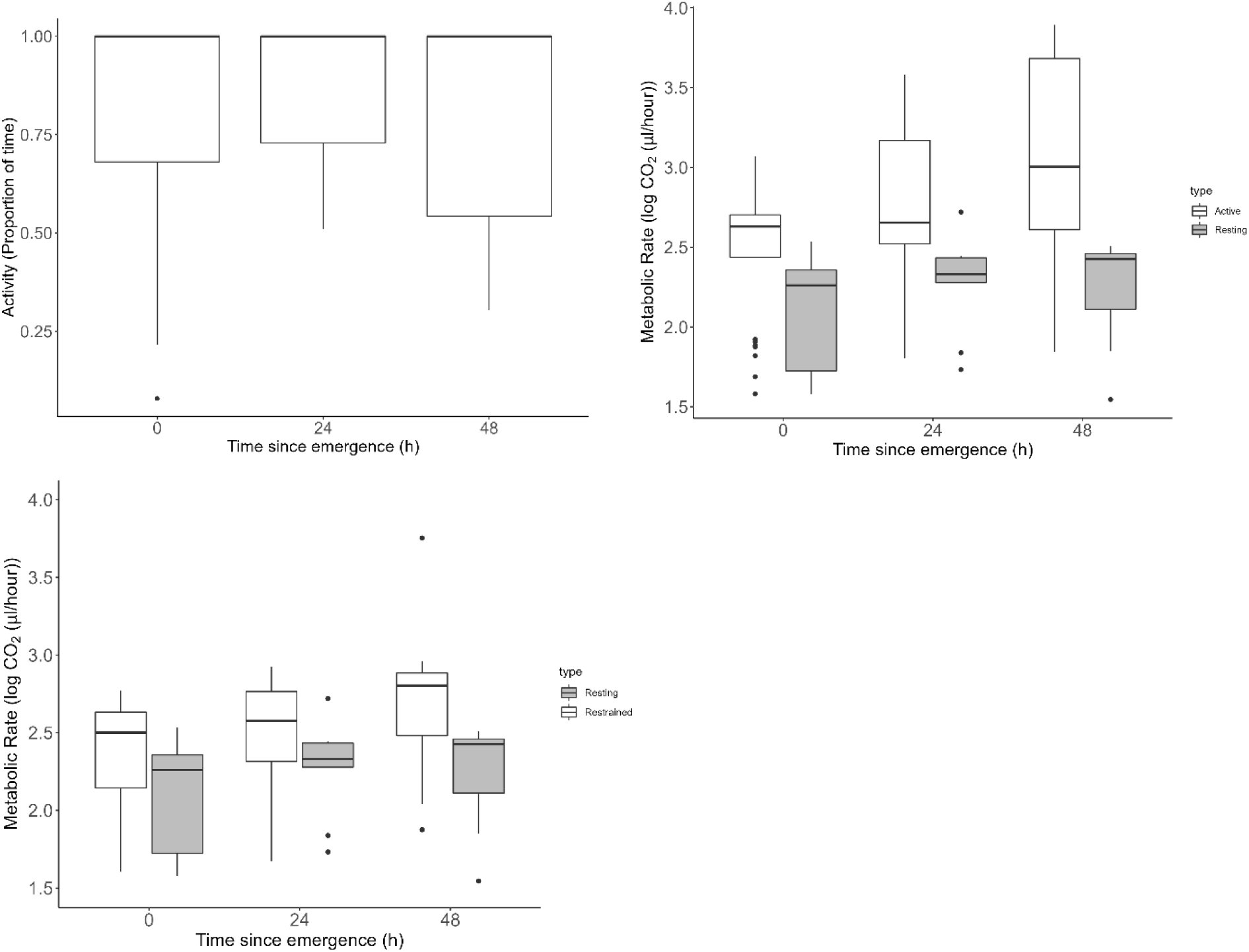
a) Proportion of time bees spent active during unrestrained metabolic recordings on the day of eclosion (0h, n=37) and 24 (n=29) and 48 hours (n=16) post-eclosion **b)** metabolic rates of unrestrained bees while resting (grey bars, 0h=18, 24h=10, 48h=7) and engaged in active movement (white bars, 0h=37, 24h=29, 48h=16) and **c**) metabolic rates of the same individual bees’ during rest while unrestrained (grey bars, 0h=37, 24h=29, 48h=17) or restrained (white bars, 0h=37, 24h=31, 48h=17).

### Both active and resting metabolic rates increase in the days post-eclosion-done

Comparing the same individuals during periods of activity versus rest, we found a significant interaction between the type of metabolic rate (RMR, AMR) and the time post-eclosion (Fig. 2b, Table S2, LMM χ^2^=7.913, df= 2, p=0.019), with AMR increasing significantly between each recording timepoint (estimated marginal means ± (emm) standard error (SE); 0-24h = -0.785±0.156, p<0.001; 24-48h - 1.337±0.254, p<0.001; 24-48h) and RMR only increasing between 0-24 hours (emm±SE -0.623±0.190, p=0.004) and 24-48h (emm± SE -0.973±0.275, p=0.002), whereas between 0-48h there was no difference (emm± SE -0.350±0.157, p=0.073). At all time points, the active metabolic rate was significantly higher than resting metabolic rate (emm±SE 0h=0.210±0.073, p=0.005; 24h=0.372±0.093, p<0.001, 48h= 0.574±0.113, p<0.001).

### Restricting movement elevates bees’ metabolic rate

Given the rarity with which unrestrained bees were stationary permitting measurement of the RMR, we measured the metabolic rates of the same individuals whilst restrained (see Methods). There was a significant interaction between the number of hours post-eclosion and the means of obtaining a metabolic rate from stationary bees (Fig. 2c, Table S3, LMM, χ^2^= 15.169, df=4, p=0.004). While there was no significant difference at 0h (emm±SE; 0h= -0.146±0.065, p=0.066), and 24h (emm±SE; 24h= - 0.153±0.083, p=0.158), the metabolic rate of restrained bees was significantly higher when recorded 48h post-emergence (48h = -0.274±0.102, p=0.021). The metabolic rate of restrained bees were significantly lower than the metabolic rates of active bees at all time points except for the day of eclosion (emm±SE; 0h= 0.093±0.050, p=0.160; 24h= 0.247±0.056, p<0.001, 48h=0.358±0.076, p<0.01). Given the metabolic rates of restrained bees differ from those of unrestrained RMR, for all future analyses we consider only unrestrained bees.

### Body mass decreases in the days post-eclosion and influences metabolic rate

Body mass decreased significantly in the hours following eclosion (Fig. 3a; Table S4, LMM, χ^2^= 2342.2, df= 2, p<0.001), between 0-24 hours (emm±SE = 0.118±0.003 p<0.001) and 24-48 hours (emm±SE = 0.068±0.004 p<0.001). Body mass also had a significant effect on metabolic rate (LMM, χ^2^= 7.650, df= 1, p=0.006).

**Figure 3.**
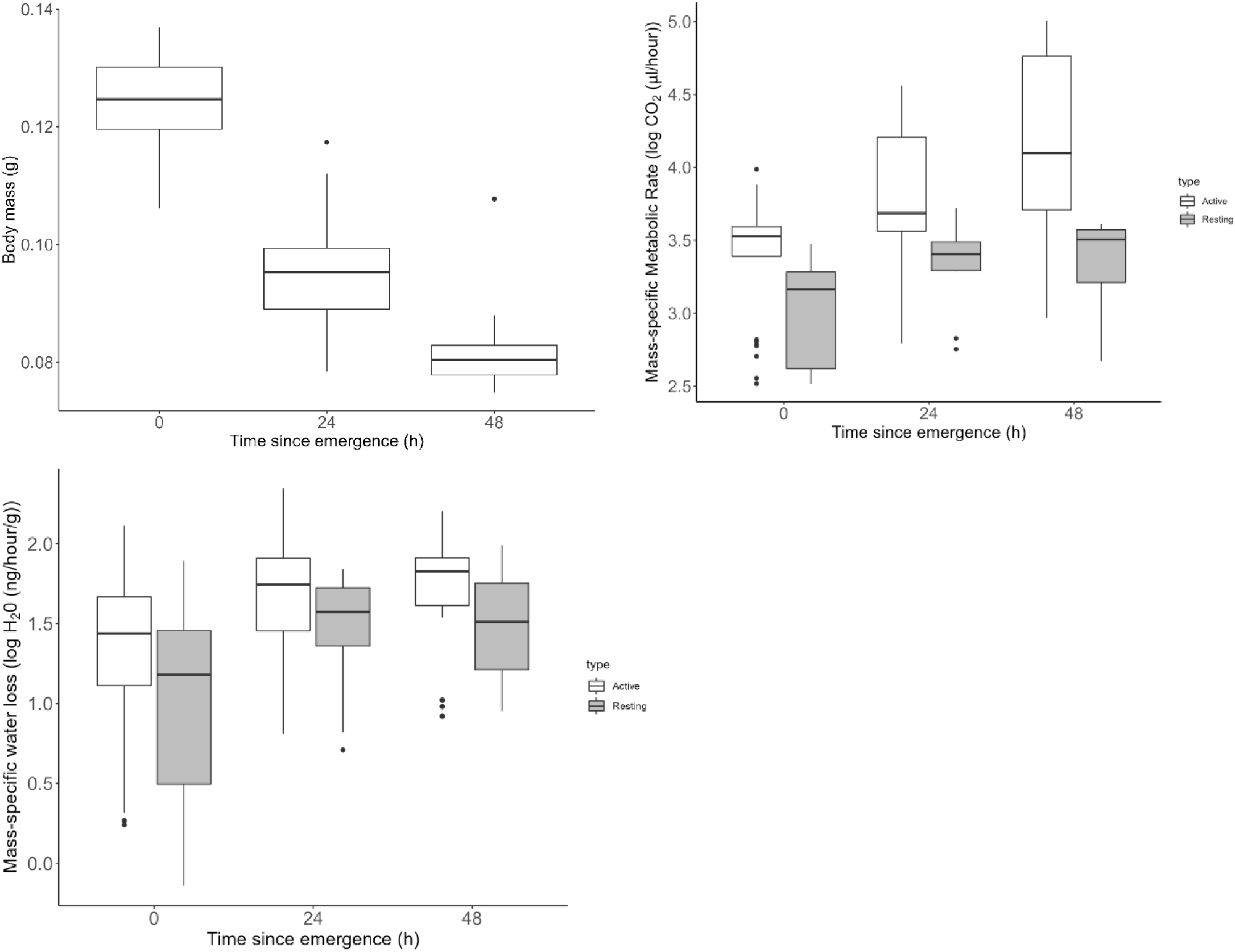
Changes in **a)** body mass **b)** mass-specific metabolic rate (metabolic rate/body mass) and **c)** mass-specific water loss of active (white bars, 0h=37, 24h=29, 48h=16) and resting bees (grey bars, 0h=18, 24h=10, 48h=7) recorded on the day of eclosion and 24 and 48 hours later.

### Mass-specific metabolic rate and water loss is highest during activity

The mass-specific metabolic rate of bees was higher when active compared to resting and the changes post-eclosion again differed between active and resting mass-specific metabolic rates (Fig. 3b, Table S5, LMM, χ^2^=7.777, df= 2, p=0.020). While the mass-specific metabolic rates of bees during activity increased significantly between each time point post-eclosion (emm ±SE 0-24h = -0.718 ± 0.175 p<0.001; 24-48h = -0.478 ±0.109, p<0.001; 0-48h = -1.196-± 0.252 p<0.001), the mass-specific metabolic rates of bees during rest only significantly increased between 0-48 hours (emm ±SE 0-48h = -0.325 ± 0.123 p=0.026) and did not change between 0-24h (−0.215 ± 0.111 p=0.138) or 24-48h hours (−0.110 ± 0.137, p=0.702). There was also a significant effect of body mass on both mass-specific RMR and AMR (LMM, χ^2^=4.269, df= 1, p=0.039), suggesting there may be a non-linear relationship between mass and metabolic rate for newly emerged bees.

Mass-specific water loss was also significantly higher during activity compared to rest (Fig. 3c, Table S6, emm±SE 0.193±0.045, p<0.001) and we observed a significant quadratic effect of time on water loss (LMM, χ^2^=6.754, df= 1, p=0.009) consistent across both activity and rest. The rate of water loss increased between 0-24 hours post-eclosion (Mean±SD 0h= 6.96± 5.44; 24 h = 12.8± 7.90 ng H_2_O·h^−1^·g^−1^·cm^−1·^s^−1^) but then stabilised between 24-48h (Mean±SD 48h= 13.5± 7.20 ng H_2_O·h^−1^·g−^1·^cm^−1^·s^−1^).

### Mass-specific metabolic scope changes significantly in the first 24 hours post-eclosion

Metabolic scope did not change with time post-emergence (AMR/RMR) (Fig. 4a, Table S7, χ^2^=1.626, df=2, p=0.443), whereas mass-specific metabolic scope did change over time (Fig. 4b, Table S8, χ^2^=31.519, df=2, p<0.001), increasing significantly between 0-24 h (emm ±SE 0-24h = -0.273 ± 0.066 p<0.001) and plateauing between 24-48 hours (24-48h= -0.134 ± 0.084 p=0.256; 0-48h= -0.407 ± 0.082, p<0.001).

**Figure 4.**
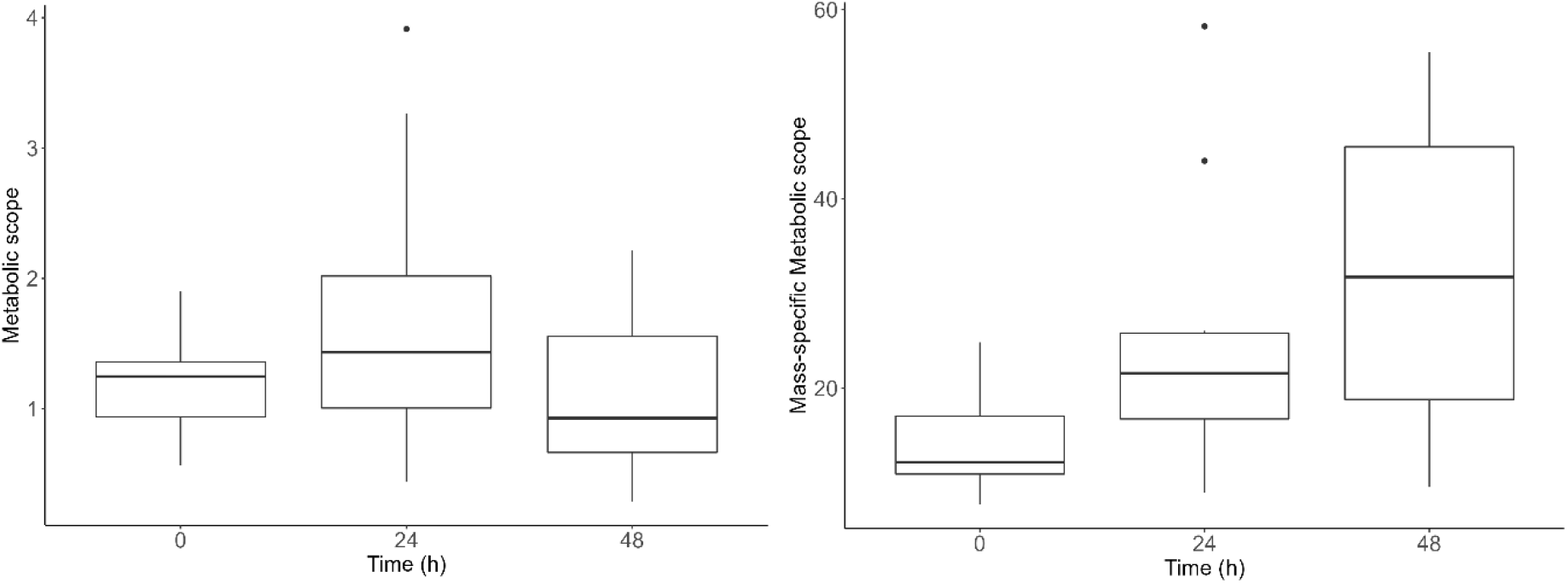
**a)** Changes in metabolic scope (active metabolic rate/resting metabolic rate) and **b)** mass-specific metabolic scope ((active metabolic rate/resting metabolic rate)/body mass) of bees recorded on the day of eclosion (n=37) and 24 (n=29) and 48 hours (n=16) later.

### Speed- and mass-specific metabolic rate and metabolic scope increase significantly in the first 48 hours post-eclosion

There was a significant quadratic effect of time on bees’ average movement speed (Fig. S2, Table S9, χ^2^=4.130, df=1, p=0.042), which increased between 0-24 hours post-eclosion (Mean ±SD 0h= 0.198 ±0.079 cm/s; 24h= 0.318± 0.181 cm/s) and then decreased again at 48 hours (48h= 0.293± 0.209 cm/s) to an average speed somewhere between that observed on the day of eclosion and 24-hours later. There was also a significant effect of time on the combined speed- and mass-specific AMR (AMR / (Body mass * average speed)) (Fig. 5a, Table, S10; χ^2^=88.409, df=2, p<0.001) which increased significantly between 0-24 hours (emm±SE 0 vs 24h= -0.199 ± 0.061, p=0.006) and 24-48 hours (24 vs 48h = -0.449 ± 0.074, p<0.001; 0 vs 48h = -0.449 ± 0.074, p<0.001).

**Figure 5.**
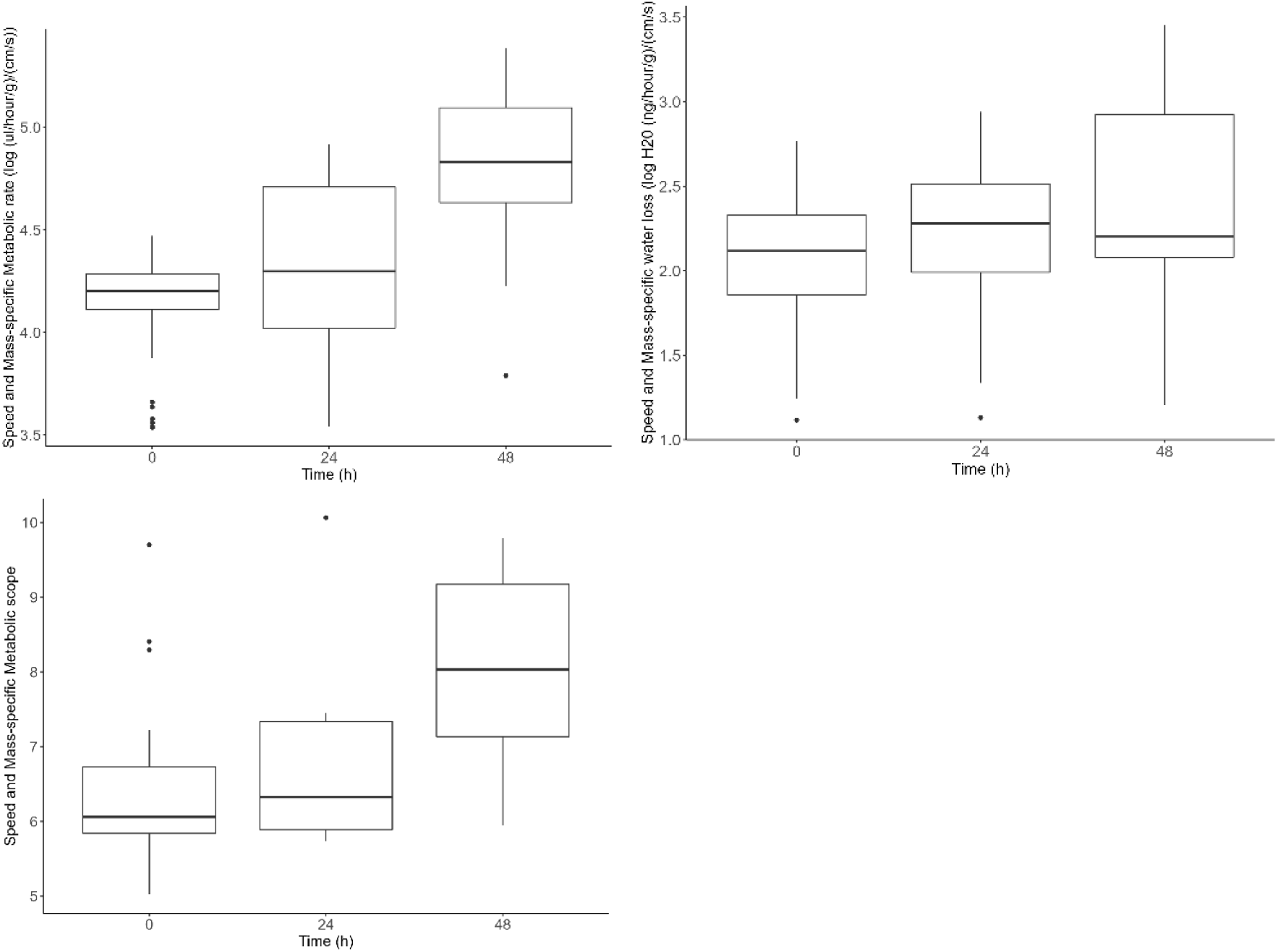
Speed- and mass-specific **a)** metabolic rate **b)** water loss and **c)** scope of actively moving bees recorded on the day of eclosion (n=37) and 24 (n=29) and 48 hours (n=16) later.

There was no significant change in speed- and mass-specific water loss (Water loss / (Body mass * Average speed)) (Fig. 5b, Table S11, χ^2^=5.545, df= 2, p=0.062) with time, however, speed- and mass-specific scope ((AMR/RMR) / (Body mass * average speed)) changed significantly with time post-emergence (Fig. 5c, Table S12, χ^2^=8.279, df= 2, p=0.016), increasing significantly between 0-48 hours (emm±SE 0 vs 48h = -0.670 ± 0.247, p=0.034; 0 vs 24h = -0.158 ± 0.220, p=0.755; 24 vs 48h = -0.512 ± 0.280, p=0.179)

## Discussion

Holometabolous insects undergo considerable behavioural and physiological changes in the hours and days post-eclosion as adults (reviewed in Chapman 2013). A limited number of studies have examined changes in insect metabolic rates during this period, however, most have focussed on resting metabolic rates (Terblanche et al. 2004; Piiroinen et al 2010) and there have been few examinations of changes in active metabolic rate and metabolic scope immediately post-eclosion (but see Schiper et al. 2009). Here, by combining flow-through respirometry with automated behaviour tracking, we show that the physiological shifts associated with eclosion are reflected in significant changes in RMR, AMR and mass-specific metabolic scope of newly emerged honeybees, providing new insights into the metabolic dynamics experienced by holometabolous insects in the initial days post-eclosion.

As has frequently been shown in insects, metabolic rates of bees engaged in activity were significantly higher than at rest (Heinrich, 1975; Josephson, 1975; Lighton, 1989; Niven & Scharlemann, 2005). Both AMR and RMR increased significantly post-eclosion, reflecting similar findings for RMR in beetles (Piiroinen et al., 2010) and RMR (Terblanche et al., 2004) and AMR in flies (Melvin, Van Voorhies & Ballard 2007). While AMR increased between all time points (from 0.39 to 2.37 ml h^-1^), RMR increased in the first 24 hours (from 0.16 to 0.23 ml h^-1^), before decreasing again (to 0.21 ml h^-1^). Mass-specific AMR and RMR also both increased significantly over 48 hours (AMR from 3.22 to 29.26, RMR from 1.33 to 2.73 ml h^-1^g^-1^) while body mass declined. The increase in metabolic rate is likely driven by the physiological changes that occur in the days immediately following eclosion, such as hypopharyngeal gland development (Crailsheim & Stolberg, 1989; Huang & Robinson, 1996), increases in juvenile hormone production (Jassim et al., 2000; Sullivan et al., 2003; Wegener et al., 2009) and changes in lipid metabolism (Toth & Robinson, 2005). Comparing the flight metabolic rates of one and three-day old bees, Schipper et al. (2009) also observed a 68% increase in metabolic rate (from *ca*. 4 to 7 ml h^-1^) that they found coincided with observed structural changes in flight muscles over the same period. However, other studies that have compared the metabolic rates of in-hive bees over longer time frames (Kovac et al., 2007), or between in-hive bees and older foraging bees (Stabentheiner et al., 2003), have observed no changes in RMR. We found that following the rapid initial increase between 0-24 hours that RMR began to decline again, suggesting such increases in RMR are transient. Indeed, the U-shaped relationship we observed for mass-specific water loss over time may reflect this. Bees experienced a rapid increase in mass-specific water loss in the first 24 hours, which also decreased by 48 hours post-eclosion. High levels of water loss are typical in newly emerged insects (Chapman 2013) and likely contribute to the decrease in body mass we observed over the first 48 hours. Complete cuticle sclerotization takes some time post-eclosion and in the interim insects are vulnerable to water loss (Gibbs & Markow, 2001; Hadley 1994). Over this same period insects also experience hormonal changes that are known to affect water balance (Nijhout 2021). It is likely that as the bees’ cuticle matured and hardened, and their hormone levels stabilised, their ability to retain water improved, which may have driven the observed decrease in water loss after 24 hours.

To our knowledge, the only previous study that has measured the metabolic rates of newly eclosed honeybees at rest used restrained bees (Allen 1959). When we compared the metabolic rates of resting bees that were either restrained or unrestrained, we found the metabolic rates of restrained bees to be significantly elevated, likely due to the stress associated with confinement, reflecting similar findings in other insects (Terblanche et al. 2004; Perl & Niven 2018). Therefore, we caution against the use of restraint in future studies of insect resting metabolic rates. In our study, newly emerged bees spent more than two thirds of their time active, and this was consistent over the first 48 hours. As highlighted by Harrison & Fewell (2002), these high activity levels are one of the major challenges for measuring the RMR of free moving bees, and one we sought to address by using the open access, automated behaviour tracking software ‘TRex’. The software permitted us to measure both resting and active metabolic rate in the *same* individuals over three different time points, thereby permitting us to examine the ontogeny of intra-specific variation in metabolic scope (defined here as the difference between an organisms’ resting and active metabolic rate while walking). Another benefit of using automated behaviour tracking is that rather than categorising the activity of the bees into different levels by hand using an ethogram (e.g. stationary, slow walking, fast walking), we could extract the exact movement speed of bees and, consequently, examine changes in speed- and mass-specific metabolic rate, water loss and scope over time.

Several studies have shown that an animal’s metabolic scope increases with the onset of specific behaviours, such as mating or flight, and declines again as an organism begins to senesce (Clark et al. 2013; Cucco et al. 2012; Javal et al. 2019; Smit et al. 2021; Schippers et al. 2009). Here we observed a quadratic relationship between time and bees’ movement speed, that increased between 0-24 hours before declining again between 24-48h to a speed between that observed at 0 and 24 hours. Speed- and mass-specific metabolic rate increased between each time point (from 15.42 at 0h to 93.98 ml h^-1^g^-1^ cm s^−1^at 48 h), whereas speed and mass-specific scope only increased after 48 hours (from 1.79 at 0h to 6.31 at 48h). There was no change in speed-and mass-specific water loss over time. While other studies have reported an increase in both activity and metabolic rate in the days post-eclosion, for example in the long horn beetle (*Cacosceles newmannii*) (Arnold et al. 2016), others have observed an increase in active metabolic rate without an associated change in movement speed, for example in the fruit fly *Drosophila melanogaster* (Melvin, van Voorhies & Ballard 2007). Javal *et al*. (2019) reported an increase in mass-specific aerobic scope from 1.68 in larvae to 5.67 ml h^-1^g^-1^ in adult longhorn beetles (*Cacosceles newmannii*), however to our knowledge no other studies have examined changes in aerobic scope or speed-specific metabolic rates in the same individuals, in the immediate days post-eclosion.

The activity and metabolic measures we have observed here suggest that in addition to the activity-driven changes in metabolic scope observed between developmental stages (Javal et al. 2019), or over longer periods post-eclosion (Schippers et al. 2009), there are also shifts in metabolism within the first 48 hours, some of which may be transient. These rapid changes could be driven by some of the maturation processes outlined previously, such as hormonal changes and flight muscle development and may reflect an initial period of flux prior to these physiological processes reaching equilibrium (Schmidt-Nielsen, 1984). The initial increase in movement speed may also reflect exploratory behaviour seen in other insects immediately following eclosion (Sokolowski 1985; Toth et al. 2005; Kirkton & Harrison 2006; Arnold et al. 2016). Another driver of the initial increase in movement speed and speed-specific metabolic rate and scope may be changes in the energy source used to fuel behaviour. As with many insects, very early post-eclosion bees rely on stored energy reserves such as lipids accumulated during the larval stage (Ziegler & Schultz 1986). Bees were only fed the artificial protein and carbohydrate-rich diet following the 0h recording, so the initial peak in movement speed and other metabolic measures observed in the recording at 24 hours post-eclosion may result from their adjustment to this new dietary source. However, in this study we only recorded CO_2_ emissions; testing for shifts in respiratory substrate would require recording of O_2_ consumption as well in order to calculate the respiratory quotient (RQ), the ratio of CO_2_ production to O_2_ consumption which provides an indication of which macronutrients are being metabolised (Lighton 2008).

These challenges of balancing increasing activity levels with the regulation of water exchange and energy reserves may have contributed to the high mortality that we observed in bees beyond 48 hours post-eclosion, suggesting that this period may be critical for survival and physiological stabilization in honeybee workers. Others have also found newly emerged bees to be at high risk of mortality and suggest this may be attributable to the energetic costs associated with metabolic adjustment in these first few days (Toth & Robinson, 2007; Harrison et al. 2012). Our method of bee husbandry may also have contributed to the high mortality, for example the diet provided may not have been sufficient to meet the shifting metabolic needs, such as the changes in lipid metabolism experienced by bees in the first few days post-eclosion (Schipers et al. 2006). Pollen is bees’ main source of lipids, and the artificial diet was lacking this. Furthermore, housing these social insects in isolation, while it enabled us to track individuals without marking them, which can itself induce stress, may have also induced higher mortality due to reduced social immunity (Ugelvig & Cremer 2007), the prohibition of food sharing (Goins & Sukumar 2013) and an overall increase in stress (Korb & Schneider 2007; Lihoreau et al. 2009).

This study provides insights into the physiology and behaviour of newly emerged honeybee workers. The high mortality rate beyond 48 hours, coupled with significant changes in metabolic rates, body mass, water loss and speed-and mass-specific metabolic scope, underscores the importance of the early post-eclosion period for survival and metabolic stabilization in bees. Our findings align with earlier research on insect metabolic rates, water regulation, and energy demands during early adult maturation. Our novel approach of combining precise activity and movement speed measurements with flow-through respirometry, goes a step further, permitting us to calculate speed- and mass-specific metabolic rates, to directly tie metabolic output to behaviour, providing a valuable tool to capture the dynamic energy costs of animal movement. Future studies should continue exploring the mechanisms driving intra-individual metabolic changes and investigate how environmental stressors and developmental conditions impact the survival of honeybee workers and ultimately colony functioning.

## Contributions

EN, JN and GV designed the study. MRA designed and built the experimental set up for behaviour tracking during flow-through respirometry and wrote the code for extracting the data on movement speed. GV collected the data. EN and GV analysed the data. GV wrote the first draft of the manuscript. All authors commented on the final version of the manuscript.

## Acknowledgements

This work is funded via a UKRI FLF awarded to EN (MR/T021691/1) and a C B Dennis Trust grant awarded to EN and JN. GV was supported by an Erasmus+ studentship and MRA by a Leverhulme Trust-funded studentship.

## Data availability

All data and scripts used in this study are open access and have been uploaded alongside this manuscript.

