## Supplementary Materials for "Shifts in honeybee worker metabolism immediately post-eclosion"

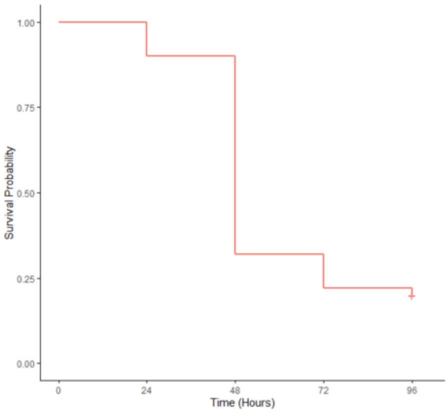


**Figure S1** Kaplan-Meier survival curve representing honeybee survival probability over time. The x-axis represents time in hours from the time of emergence (0 hours) to 96 hours post-emergence. The solid red line represents the survival curve, indicating the proportion of bees recorded on the day of emergence (n = 37) surviving for re-recording at each time point.

| | Activity | | |
|---|---|---|---|
| *Predictors* | *Estimates* | *CI* | *p* |
| (Intercept) | 5.90 | 0.00 – Inf | **<0.001** |
| time1 | 0.90 | 0.00 – Inf | 0.553 |
| time2 | 1.15 | 0.00 – Inf | 0.423 |
| **Random Effects** | | | |
| $\sigma^2$ | 0.07 | | |
| $\tau_{00}$ bee_id | 0.00 | | |
| N bee_id | 37 | | |
| Observations | 82 | | |
| Marginal $R^2$ / Conditional $R^2$ | 0.141 / NA | | |

**Table S1:** Model summary for comparison of activity over time post-eclosion. Model formula used [activity ~ time + (1|bee_id), family = beta_family(link = "logit")].

| | Metabolic Rate | | | |
|---|---|---|---|---|
| *Predictors* | *Estimates* | *CI* | *Statistic* | *p* |
| (Intercept) | 6.98 | -Inf – Inf | 5.34 | **<0.001** |
| 24 hours | -0.62 | -Inf – Inf | -4.51 | **<0.001** |
| 48 hours | 0.08 | -Inf – Inf | 1.86 | 0.065 |
| Active | 0.19 | -Inf – Inf | 6.99 | **<0.001** |
| log10(body_mass) | 4.40 | -Inf – Inf | 3.41 | **0.001** |
| 24 hours:Active | -0.09 | -Inf – Inf | -2.58 | **0.011** |
| 48 hours:Active | -0.01 | -Inf – Inf | -0.17 | 0.863 |
| **Random Effects** | | | | |
| $\sigma^2$ | 0.06 | | | |
| $\tau_{00\ bee\_id}$ | 0.16 | | | |
| ICC | 0.74 | | | |
| $N_{bee\_id}$ | 37 | | | |
| Observations | 117 | | | |
| Marginal $R^2$ / Conditional $R^2$ | 0.247 / 0.804 | | | |

**Table S2:** Model summary for comparison of active and resting metabolic rates over time post-eclosion. Model formula used [log10(metabolic rate) ~ time + activity_type + log10(body_mass) + time*activity_type + (1| bee_id)].

| | Metabolic Rate | | | |
|---|---|---|---|---|
| *Predictors* | *Estimates* | *CI* | *Statistic* | *p* |
| (Intercept) | 5.08 | -Inf – Inf | 6.35 | **<0.001** |
| 24 hours | -0.41 | -Inf – Inf | -4.89 | **<0.001** |
| 48 hours | 0.05 | -Inf – Inf | 1.72 | 0.087 |
| Active | 0.22 | -Inf – Inf | 9.34 | **<0.001** |
| Resting | -0.20 | -Inf – Inf | -6.71 | **<0.001** |
| log10(body_mass) | 2.55 | -Inf – Inf | 3.22 | **0.002** |
| 24 hours:Active | -0.11 | -Inf – Inf | -3.63 | **<0.001** |
| 48 hours:Active | -0.00 | -Inf – Inf | -0.10 | 0.917 |
| 24 hours:Resting | 0.08 | -Inf – Inf | 2.03 | **0.043** |
| 48 hours:Resting | 0.02 | -Inf – Inf | 0.49 | 0.625 |
| **Random Effects** | | | | |
| $\sigma^2$ | 0.05 | | | |
| $\tau_{00\ bee\_id}$ | 0.13 | | | |
| ICC | 0.73 | | | |
| N $_{bee\_id}$ | 37 | | | |
| Observations | 202 | | | |
| Marginal $R^2$ / Conditional $R^2$ | 0.205 / 0.786 | | | |

**Table S3:** Model summary for comparison of active, resting and restrained bees' metabolic rates over time post-eclosion. Model formula used [log10(metabolic rate) ~ time + activity_type + log10(body_mass) + time*activity_type + (1| bee_id)]

| | Body Mass | | |
|---|---|---|---|
| *Predictors* | *Estimates* | *CI* | *p* |
| (Intercept) | -1.01 | -Inf – Inf | **<0.001** |
| 24 hours | 0.10 | -Inf – Inf | **<0.001** |
| 48 hours | -0.02 | -Inf – Inf | **<0.001** |
| **Random Effects** | | | |
| $\sigma^2$ | 0.00 | | |
| $\tau_{00\ bee\_id}$ | 0.00 | | |
| ICC | 0.61 | | |
| $N_{bee\_id}$ | 37 | | |
| Observations | 249 | | |
| Marginal $R^2$ / Conditional $R^2$ | 0.843 / 0.938 | | |

**Table S4:** Model summary for changes in body mass over time post-eclosion. Model formula used [log10(body_mass) ~ time + (1| bee_id)]

| | Mass-specific Metabolic Rate | | |
|---|---|---|---|
| *Predictors* | *Estimates* | *CI* | *p* |
| (Intercept) | 3.54 | -Inf – Inf | **<0.001** |
| 24 hours | -0.27 | -Inf – Inf | **<0.001** |
| 48 hours | 0.03 | -Inf – Inf | 0.517 |
| Active | 0.21 | -Inf – Inf | **<0.001** |
| 24 hours * Active | -0.09 | -Inf – Inf | **0.012** |
| 48 hours * Active | -0.01 | -Inf – Inf | 0.861 |
| **Random Effects** | | | |
| $\sigma^2$ | 0.06 | | |
| $\tau_{00\ bee\_id}$ | 0.13 | | |
| ICC | 0.67 | | |
| $N_{bee\_id}$ | 37 | | |
| Observations | 117 | | |
| Marginal $R^2$ / Conditional $R^2$ | 0.315 / 0.772 | | |

**Table S5** Model summary for comparison of active and resting mass-specific metabolic rate over time post-eclosion. Model formula used [log10(ms_co2)~time+type+time:type+(1|bee_id)]

| | Mass-specific Water Loss | | |
|---|---|---|---|
| *Predictors* | *Estimates* | *CI* | *p* |
| (Intercept) | 1.23 | -Inf – Inf | **<0.001** |
| time | 0.37 | -Inf – Inf | **<0.001** |
| Active | 0.10 | -Inf – Inf | **<0.001** |
| time^2 | -0.12 | -Inf – Inf | **0.011** |
| **Random Effects** | | | |
| $\sigma^2$ | 0.04 | | |
| $\tau_{00}$ bee_id | 0.19 | | |
| ICC | 0.82 | | |
| N bee_id | 37 | | |
| Observations | 117 | | |
| Marginal $R^2$ / Conditional $R^2$ | 0.103 / 0.836 | | |

**Table S6** Model summary for comparison of active and resting mass-specific water loss over time post-eclosion. Model formula used [log10(mass_specific_waterloss) ~ time + I(time^2) + activity_type + log10(body_mass) + (1| bee_id)].

| | Metabolic Scope | | |
|---|---|---|---|
| *Predictors* | *Estimates* | *CI* | *p* |
| (Intercept) | 0.29 | -Inf – Inf | **<0.001** |
| 24 hours | -0.06 | -Inf – Inf | 0.214 |
| 48 hours | 0.03 | -Inf – Inf | 0.594 |
| **Random Effects** | | | |
| $\sigma^2$ | 0.03 | | |
| $\tau_{00\ bee\_id}$ | 0.01 | | |
| ICC | 0.30 | | |
| N $_{bee\_id}$ | 26 | | |
| Observations | 35 | | |
| Marginal $R^2$ / Conditional $R^2$ | 0.039 / 0.325 | | |

**Table S7:** Model summary for comparison of scope over time post-eclosion. Model formula used [log10(scope)~time+(1|bee_id)].

| | Mass-specific Metabolic Scope | | |
|---|---|---|---|
| *Predictors* | *Estimates* | *CI* | *p* |
| (Intercept) | 1.21 | -Inf – Inf | **<0.001** |
| 24 hours | -0.23 | -Inf – Inf | **<0.001** |
| 48 hours | 0.05 | -Inf – Inf | 0.279 |
| **Random Effects** | | | |
| $\sigma^2$ | 0.07 | | |
| $\tau_{00\ bee\_id}$ | 0.01 | | |
| ICC | 0.13 | | |
| N $_{bee\_id}$ | 37 | | |
| Observations | 82 | | |
| Marginal $R^2$ / Conditional $R^2$ | 0.260 / 0.355 | | |

**Table S8**: Model summary for comparison of mass-specific scope over time post-eclosion Model formula used (log10(MSScope)~time+(1|bee_id),data=df_scope)

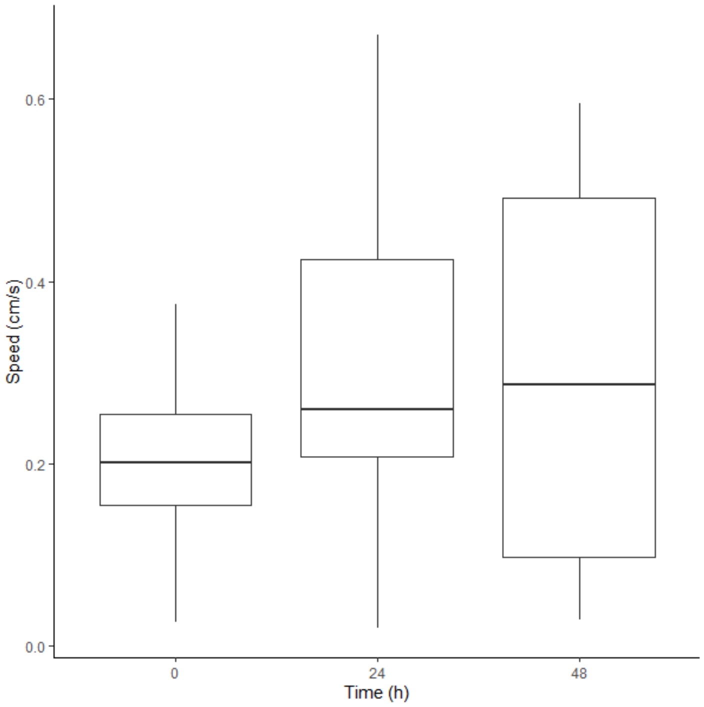


**Figure S2:** Movement speed (cm/s) of active bees at 0 (n=37), 24 (n=29) and 48 hours (n=16) post-eclosion.

| | Movement Speed | | |
|---|---|---|---|
| *Predictors* | *Estimates* | *CI* | *p* |
| (Intercept) | 0.20 | -Inf – Inf | **<0.001** |
| time | 0.01 | -Inf – Inf | **0.007** |
| time^2 | -0.00 | -Inf – Inf | **0.046** |
| **Random Effects** | | | |
| $\sigma^2$ | 0.02 | | |
| $\tau_{00\ bee\_id}$ | 0.00 | | |
| ICC | 0.20 | | |
| N $_{bee\_id}$ | 37 | | |
| Observations | 74 | | |
| Marginal $R^2$ / Conditional $R^2$ | 0.128 / 0.306 | | |

**Table S9:** Model summary for comparison of movement speed over time post-eclosion. Model formula used [movement_speed~time+I(time^2)+(1|bee_id)].

| | Speed-mass-specific Metabolic Rate | | |
|---|---|---|---|
| *Predictors* | *Estimates* | *CI* | *p* |
| (Intercept) | 4.42 | -Inf – Inf | **<0.001** |
| 24 hours | -0.28 | -Inf – Inf | **<0.001** |
| 48 hours | -0.08 | -Inf – Inf | **0.035** |
| **Random Effects** | | | |
| $\sigma^2$ | 0.04 | | |
| $\tau_{00\ bee\_id}$ | 0.08 | | |
| ICC | 0.64 | | |
| N $_{bee\_id}$ | 37 | | |
| Observations | 74 | | |
| Marginal $R^2$ / Conditional $R^2$ | 0.348 / 0.763 | | |

**Table S10:** Model summary for comparison of speed- and mass-specific metabolic rate over time post-eclosion Model formula used [log10(speed_mass_metabolic_rate)~time+(1|bee_id)].

| | Speed-mass-specific Water Loss | | |
|---|---|---|---|
| *Predictors* | *Estimates* | *CI* | *p* |
| (Intercept) | 2.20 | -Inf – Inf | **<0.001** |
| 24 hours | -0.12 | -Inf – Inf | **0.037** |
| 48 hours | -0.02 | -Inf – Inf | 0.791 |
| **Random Effects** | | | |
| $\sigma^2$ | 0.11 | | |
| $\tau_{00\ bee\_id}$ | 0.11 | | |
| ICC | 0.50 | | |
| N $_{bee\_id}$ | 37 | | |
| Observations | 74 | | |
| Marginal $R^2$ / Conditional $R^2$ | 0.043 / 0.521 | | |

**Figure S11**: Model summary for comparison of speed- and mass-specific water loss over time post-eclosion [log10(speed_mass_water)~time+(1|bee_id)].

| | Speed-mass-specific Scope | | |
|---|---|---|---|
| *Predictors* | *Estimates* | *CI* | *p* |
| (Intercept) | 3.09 | -Inf – Inf | **<0.001** |
| 24 hours | -0.28 | -Inf – Inf | **0.028** |
| 48 hours | -0.12 | -Inf – Inf | 0.399 |
| **Random Effects** | | | |
| $\sigma^2$ | 0.24 | | |
| $\tau_{00\ bee\_id}$ | 0.06 | | |
| ICC | 0.21 | | |
| N $_{bee\_id}$ | 26 | | |
| Observations | 35 | | |
| Marginal $R^2$ / Conditional $R^2$ | 0.179 / 0.348 | | |

**Figure 12:** Model summary for comparison of speed- and mass-specific scope over time post-eclosion [log10(speed_mass_scope)~time+(1|bee_id)].
